# A new chromosome-level genome assembly and annotation of *Cryptosporidium meleagridis*

**DOI:** 10.1101/2024.02.16.580748

**Authors:** Lasya R. Penumarthi, Rodrigo P. Baptista, Megan S. Beaudry, Travis C. Glenn, Jessica C. Kissinger

## Abstract

*Cryptosporidium* spp. are medically and scientifically relevant protozoan parasites that cause severe diarrheal illness in infants and immunosuppressed populations as well as animals. Although most human *Cryptosporidium* infections are caused by *C. parvum* and *C. hominis*, there are several other human-infecting species including *C. meleagridis*, which is commonly observed in developing countries. Here, we polished and annotated a long-read genome sequence assembly for *C. meleagridis* TU1867, a species which infects birds and humans. The genome sequence was generated using a combination of whole genome amplification (WGA) and long-read Oxford Nanopore Technologies sequencing. The assembly was then polished with Illumina data. The chromosome-level genome assembly is 9.2 Mbp with a contig N50 of 1.1 Mb. Annotation revealed 3,923 protein-coding genes. A BUSCO analysis indicates a completeness of 96.6% (n=446), including 430 (96.4%) single-copy and 1 (0.224%) duplicated apicomplexan conserved gene(s). The new *C. meleagridis* genome assembly is nearly gap-free and provides a valuable new resource for the *Cryptosporidium* community and future studies on evolution and host-specificity.

## Background & Summary

*Cryptosporidium* is an apicomplexan protozoan parasite of global medical, scientific, and veterinary significance that can cause moderate-to-severe diarrhea in humans and animals^1^. It is the leading cause of waterborne disease outbreaks in the US^2,3^. Though cryptosporidiosis causes illness in both immunocompromised and immunocompetent individuals, it is especially severe in immunocompromised and elderly populations as well as in children, resulting in persistent infection, malnutrition, and, in some cases, death^3-5^. In 2019, the Global Burden of Disease study found 133,422 global deaths and an annual loss of 8.2 million disability-adjusted life years (DALYs) due to *Cryptosporidium*^6^. *C. meleagridis* is an avian and mammalian-infecting *Cryptosporidium* species that was first described in turkeys^7,8^. Human *Cryptosporidium* infections are caused predominantly by *C. parvum* and *C. hominis*, but species such as *C. meleagridis* can also infect humans. In fact, *C. meleagridis* is the third most common human-infecting *Cryptosporidium* species following *C. parvum* and *C. hominis*^9^. Though generally less common, *C. meleagridis* infection has been reported to be as common as *C. parvum* in some parts of the world and can lead to death in rare cases^10,11^.

At this time, 15 of the >30 reported *Cryptosporidium* species have assembled genome sequences. However, only 8 have been annotated including *C. parvum, C. hominis*, C. *tyzzeri*, and *C. meleagridis*^12^. The latest release of the *C. meleagridis* genome strain UKMEL1 (*Cm*UKMEL1) contains gaps and is assembled into 57 contigs. Due to a highly compact genome, it is challenging to sequence the genome of *Cryptosporidium* parasites from a single individual. Since cloning of *Cryptosporidium* parasites is not possible, sequencing a small pool of individuals is preferred over bulk sequencing to reduce heterozygosity. Recently, a new method has been implemented to generate DNA sequences from *Cryptosporidium* using a whole genome amplification (WGA) approach and was tested on *C. meleagridis* strain TU1867 (*Cm*TU1867) which provided sufficient DNA for library construction and generation of a high-quality genome through long-read sequencing ^13^. Here we share a chromosome-level assembly of the *C. meleagridis* genome. The new *Cm*TU1867 genome assembly is 201,275 base pairs longer than that of *Cm*UKMEL1. The largest contig in the new assembly is 632,735 base pairs (bp) longer than the largest contig in *Cm*UKMEL1. We note a larger N50 value of 1,105,563 bp in the new *Cm*TU1867 assembly compared to *Cm*UKMEL1 which has an N50 value of 322,908 bp (Table 1). The new *Cm*TU1867 assembly and annotation provides a valuable resource to the *Cryptosporidium* community. The high-quality *C. meleagridis* genome results from a new experimental approach designed to help generate whole genome sequences from limiting amounts of genomic DNA and is an important resource that will contribute to our understanding of *Cryptosporidium* evolution and host specificity.

**Table 1.** Statistics of the *C. meleagridis and Cp*BGF genome assemblies and annotation.

| Statistics | CmTU1867 | CmUKMEL1 | CpBGF |
| --- | --- | --- | --- |
| # of contigs | 13 | 57 | 8 |
| Largest contig (bp) | 1365597 | 732862 | 1379130 |
| # T2T chromosomes | 0 | 0 | 8 |
| Total length (bp) | 9178485 | 8973200 | 9222738 |
| N50 (bp) | 1105563 | 322908 | 1108772 |
| GC (%) | 30.9 | 31.0 | 30.1 |
| # N's per 100 kbp | 0.00 | 0.00 | 0.00 |
| # of telomeres identified | 1 | 10 | 16 |
| # rRNA | 17 | 8 | 15 |
| # tRNA | 45 | 45 | 45 |
| # protein-coding genes | 3923 | 3753 | 3932 |
| Average gene length (bp) | 1824 | 1885 | 2145 |

The initial genome assembly contained 13 contigs including 8 chromosomes, 5 contigs (681-30,300 bp), 2 of which were later identified as contamination and removed (Figure 1). Two additional contigs were created manually (“contig_ 10” and “contig_11”) from the beginnings of chromosome 2 and chromosome 6 due to detection of an assembly artifact in these chromosomes. Thus, the final assembly contains 13 contigs, 8 chromosomes and contigs 9-13. Contig_9 and contig_13 have regions identical to regions of chromosome 1 and chromosome 3, respectively, but assembled separately from the full chromosomes (Table 1). One drawback to the new assembly is its lack of telomeres in comparison to *Cm*UKMEL1. We were only able to detect 1 telomere in *Cm*TU1867 on chromosome 5. Searching through the long reads, we were able to find several reads with telomeres on them that did not assemble. Though these reads did not assemble, regions of the read that did not contain the telomere pattern matched the assembly. Upon mapping these reads back to the assembly, we identified three additional telomere locations that could be placed manually (beginning and end of chromosome 3 and beginning of chromosome 4). At least 4 telomere-containing long-reads mapped to these regions with at least 1 long (>1kb) read that extended into unique regions of the chromosome. However, due to relatively low read support for these telomeres, we did not extend the ends of chromosomes in the assembly with these telomere-containing reads.

**Figure 1.**
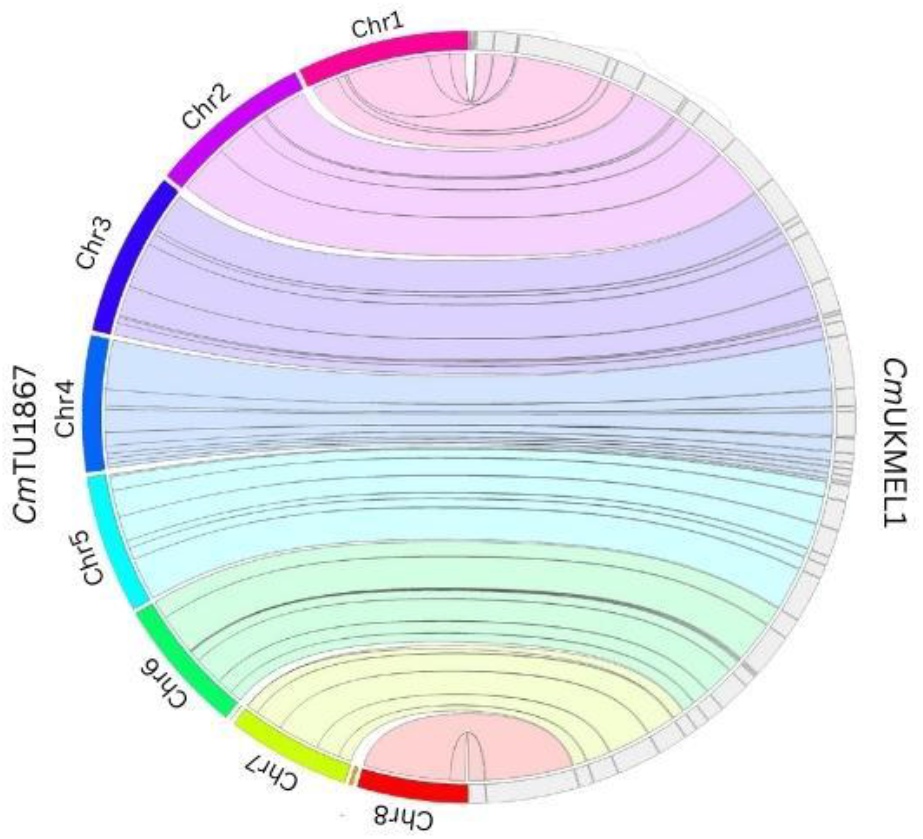
DNA synteny plot of the eight chromosome level contigs of *Cm*TU1867 (left hemisphere) and *Cm*UKMEL1 (right hemisphere). Jupiterplot between the previous *Cm*UKMEL1 genome sequence and the new *Cm*TU1867 genome sequence. Ribbons are colored with respect to the reference genome (*Cm*TU1867).

The new *Cm*TU1867 genome sequence has 10 additional ribosomal RNA genes compared to *Cm*UKMEL1. The 16 rRNA genes (excluding the 5.8S) are in clusters of 2-3 and are found on either chromosome 1, 2, 3, 7, or 8 (Figure 2). We noticed that what RNAmmer^14^ detected as 5.8S rRNAs in *Cm*TU1867 clustered separately from the 18S and the 28S rRNAs with the exception of one 5.8S (ID=cmbei_2001394) located on chromosome 2 adjacent to a 28S rRNA. Most Apicomplexans have been annotated to have the 5.8S rRNA in between the 18S and 28S rRNAs. Additional searches revealed that all but the 5.8S on chromosome 2 were 5S rRNAs. The six 5S rRNAs in *Cm*TU1867 are in 2 clusters of 3, one on chromosome 3 and the other on chromosome 7 (Figure 2). In *Cp*BGF the cluster of 5S rRNAs in chromosome 3 contains 2 rRNAs whereas in *Cp*IOWA-ATCC and *Cm*TU1867, the cluster of 5S rRNAs in chromosome 3 contains 3 rRNAs. These patterns may arise because of variation in the copy number of the 5S rRNA within a population of parasites or among different species of *Cryptosporidium* or compressions during genome assembly. When *Cm*TU1867 reads were mapped to the assembly at regions where there are 5S rRNA clusters in chromosomes 3 and 7, we saw relatively even coverage throughout the region. However, when we did this test with the *Cp*BGF reads and genome sequence, we found a 2-3x read compression at precisely the 5S regions on chromosome 3 and 7.

**Figure 2.**
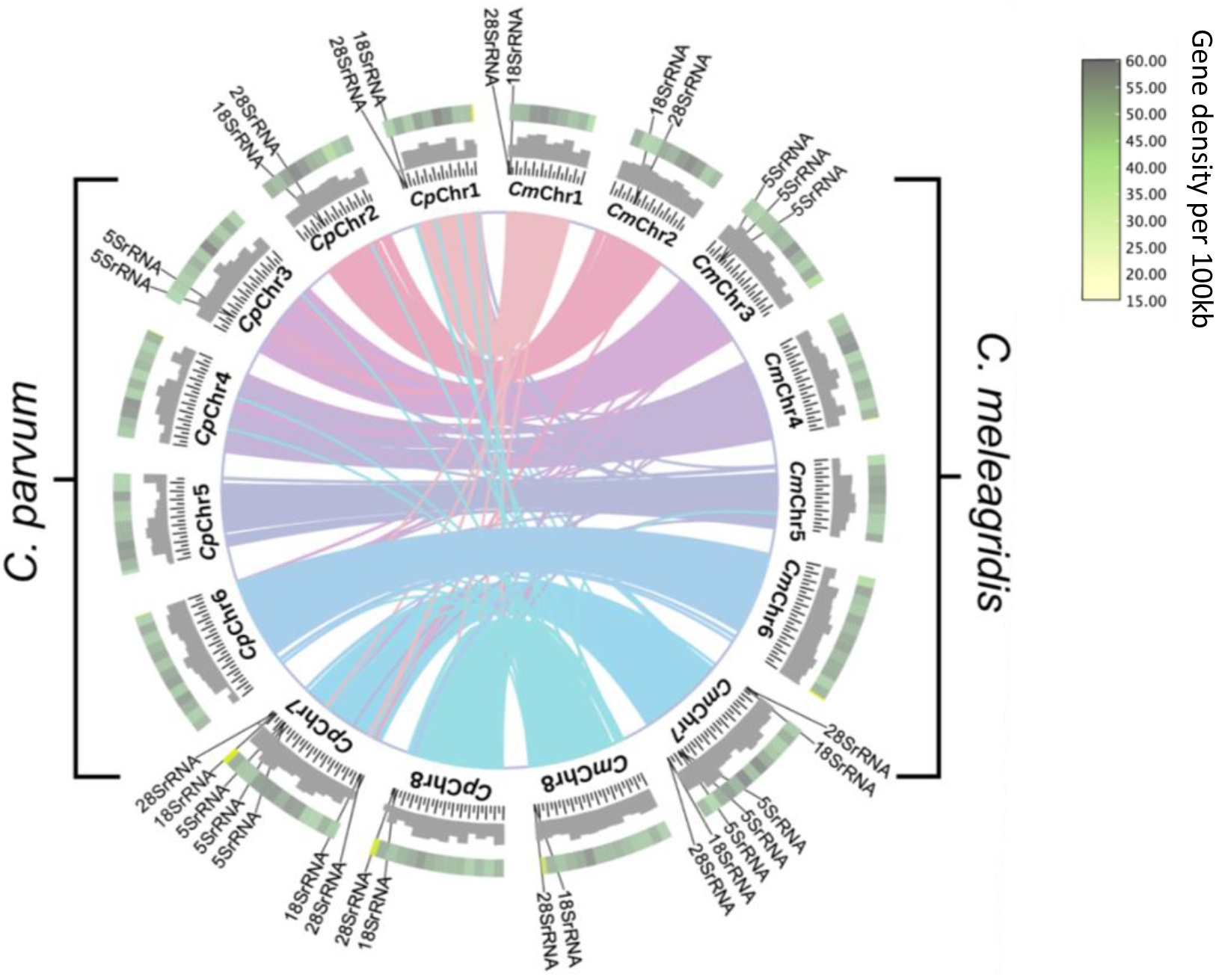
Protein synteny analysis of the eight chromosome-level contigs of *Cm*TU1967 (right hemisphere) and *Cryptosporidium parvum, Cp*BGF (left hemisphere). Circos plot rings, moving from the center to the exterior illustrate shared ortholog clusters between *Cm*TU1867 and *Cp*BGF, number of base pairs in 50,000 bp increments, GC content histogram, and gene density. Locations of rRNA genes are as indicated.

The annotation was validated by comparing the *Cm*TU1867, *Cp*BGF, and *Cm*UKMEL1 protein-coding sequences using orthology-based algorithms. We initially found a group of 5 endonucleases in *Cm*TU1867 not in *Cp*BGF or *Cm*UKMEL1. The members of this group did not have any significant hit to any *Cryptosporidium* species except *C. ubiquitum* and *C. felis* in a blastp search. However, when we ran tblastn of this gene family against annotated transcripts in CryptoDB, we found hits to *C. parvum, C. hominis, C. tyzerri, C. ryanae*, and *Cm*UKMEL1. The members of the gene family have been annotated as 18S ribosomal or non-coding RNAs in *C. parvum, C. hominis, C. tyzerri*, and *C. ryanae* and as a 5S rRNA and ncRNA in *Cm*UKMEL1 but as protein-coding genes in *C. ubiquitum*. Upon manual inspection we found that the genomic sequence for these proteins exists in the *C. parvum, C. hominis*, and *C. tyzerri* genome in varying copy number. We additionally found one putative open-reading frame (ORF) predicted by AUGUSTUS in *Cm*TU1867 Chr4 region 828518-828700 bp that was not in *Cm*UKMEL1, *Cp*BGF, or any other species according to blastp and blastn searches. We removed this putative gene from our annotations since we could not validate it with RNAseq data or evidence in any other species, but it may be a unique *C. meleagridis* gene detected by the improved assembly.

Some of the orthogroups initially detected by OrthoVenn3 fell at the ends of chromosomes in *C*.*parvum* that extended beyond the ends of the *Cm*TU1867 and *Cm*UKMEL1 chromosomes. Other times they were unannotated in one species or the other but present in the genome sequence. When we found unannotated proteins that were not initially detected by Liftoff or AUGUSTUS in *Cm*TU1867, we manually added these annotations. Ultimately, we found very few orthogroups that were unique to a species (Figure 3). A description of their investigation can be found in Table 3.

**Table 2.**
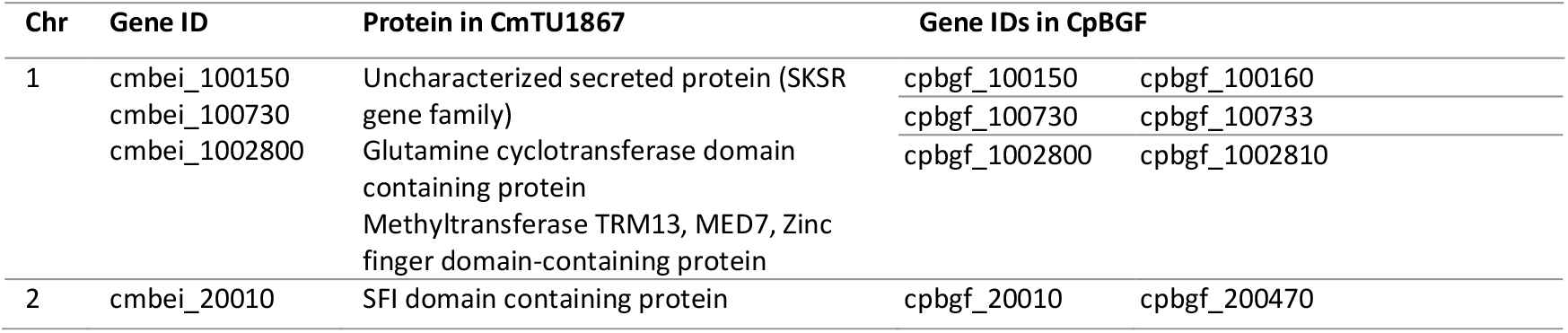
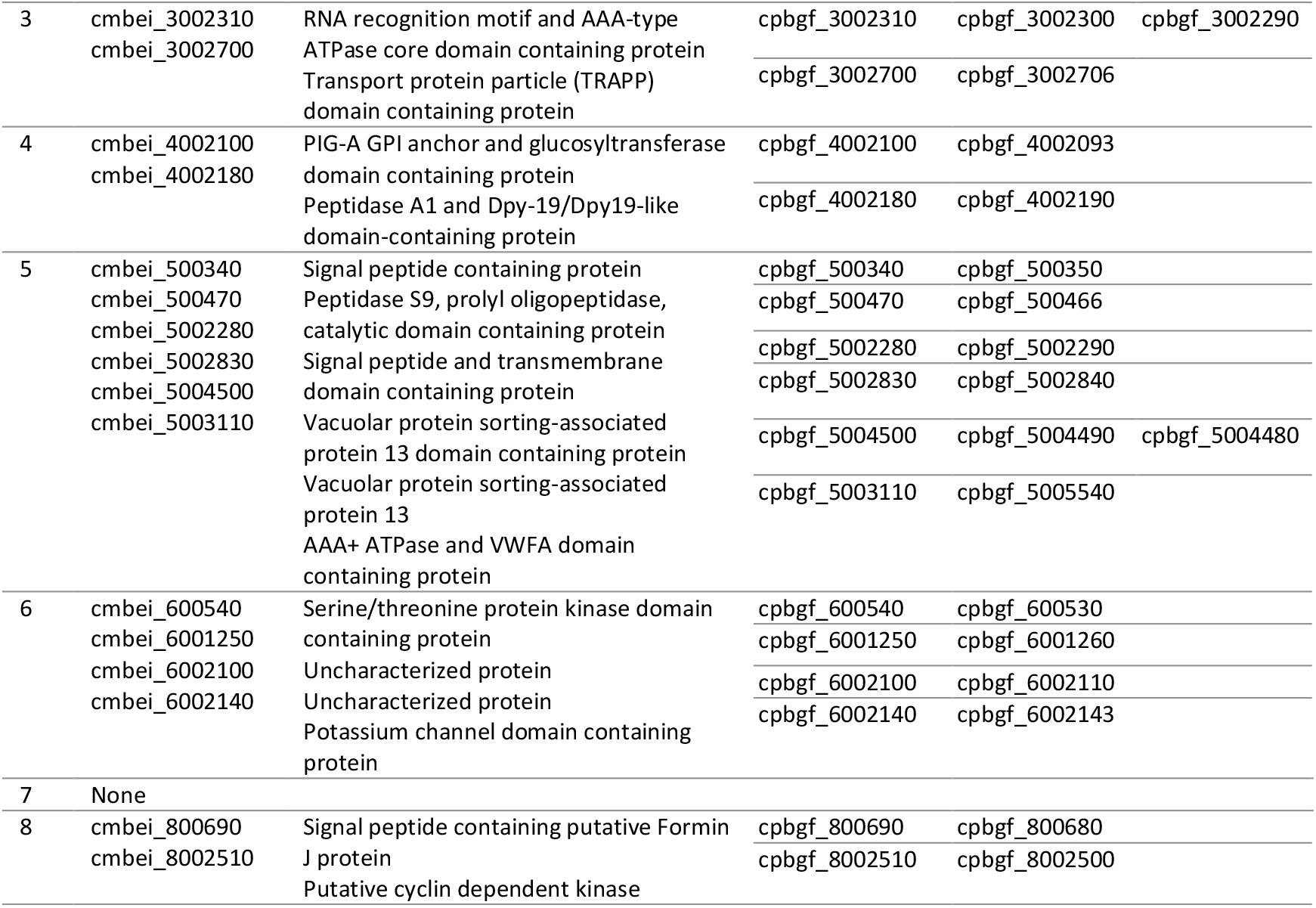
Single large, annotated genes in *Cm*TU1867 that are annotated as two or three distinct sequential genes in *Cp*BGF and/or other *Cryptosporidium* spp.

**Table 3.**
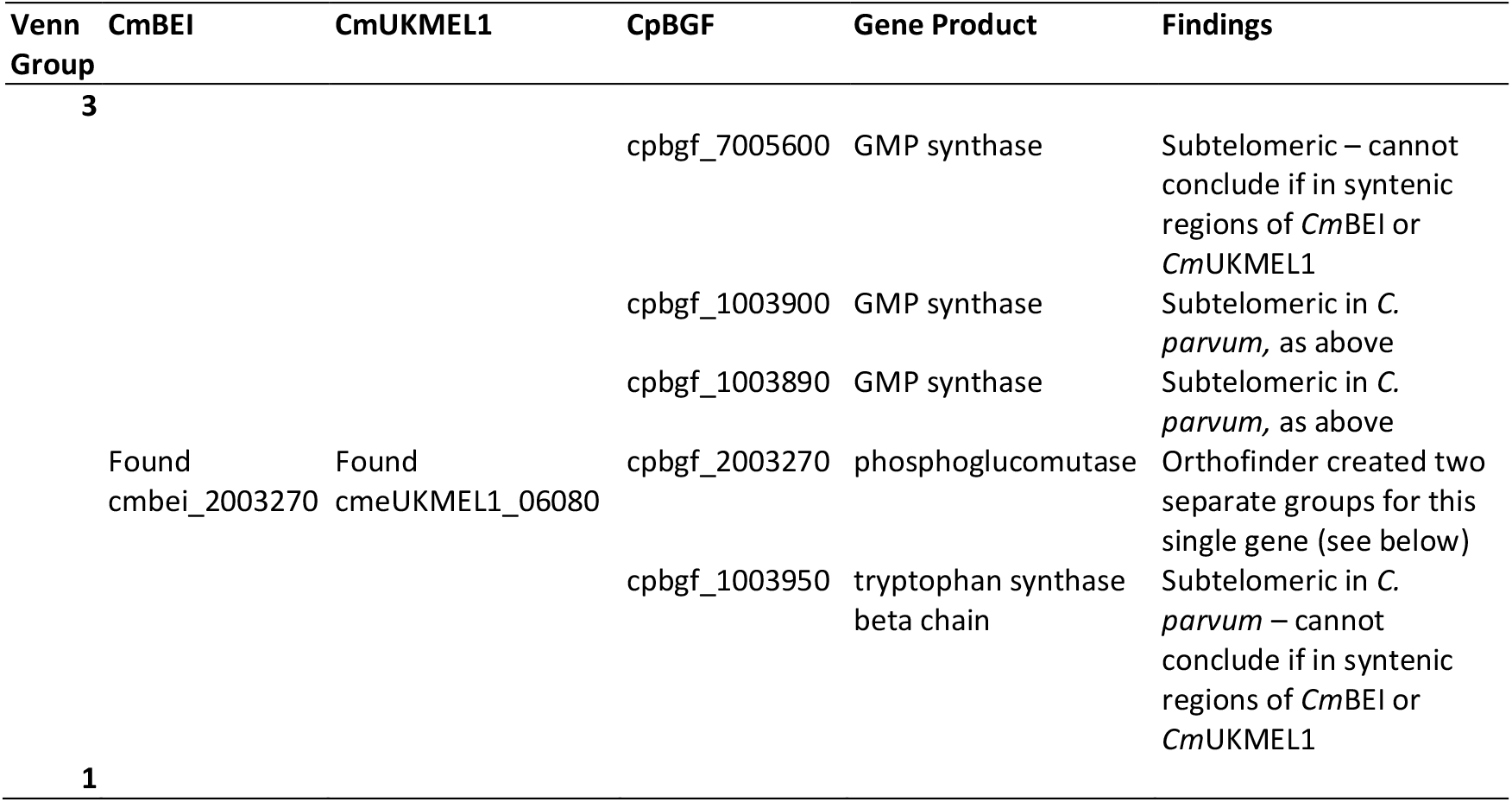
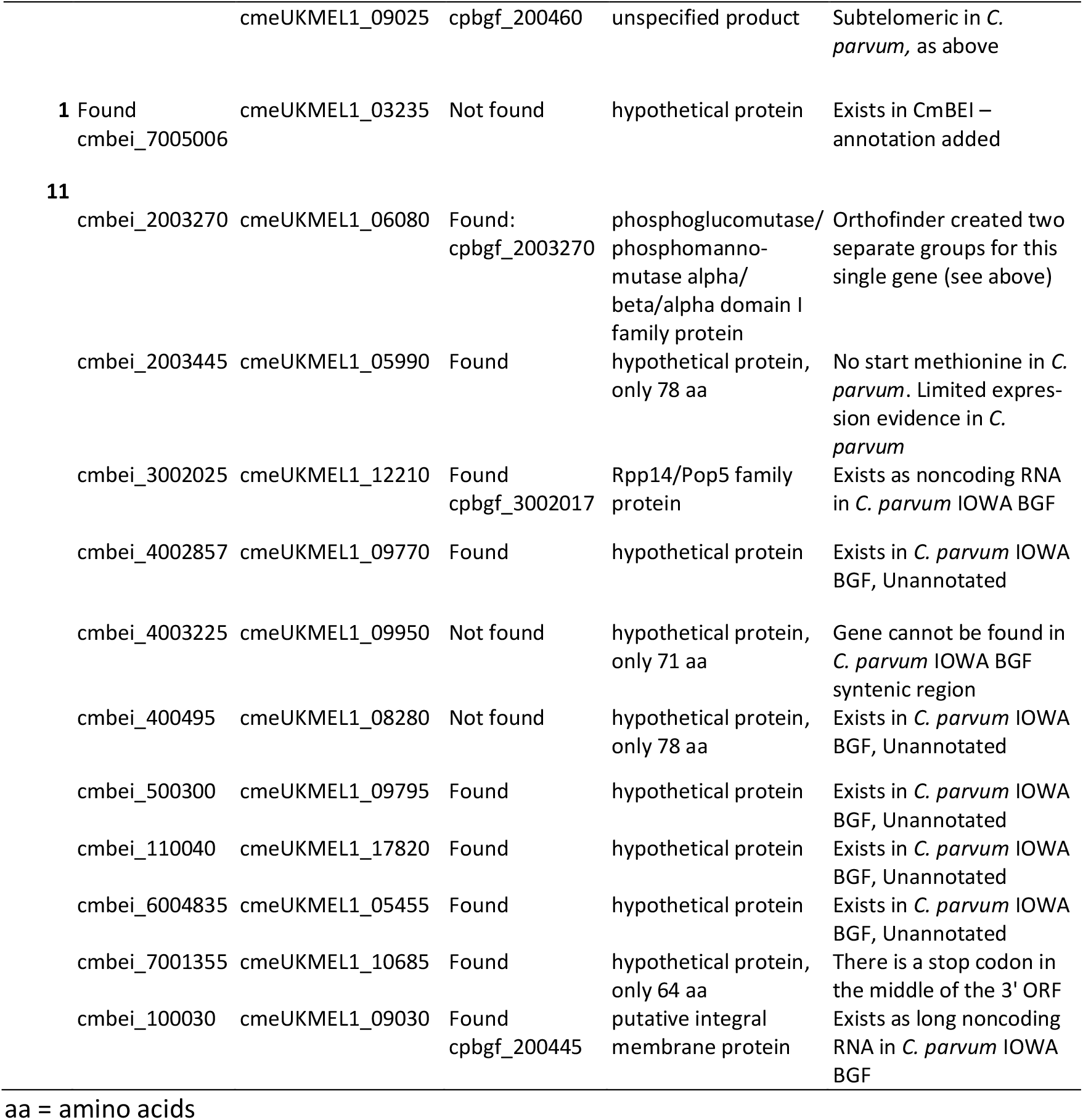
Manual validation of orthogroups not present in all species examined in Figure 5.

**Figure 3.**
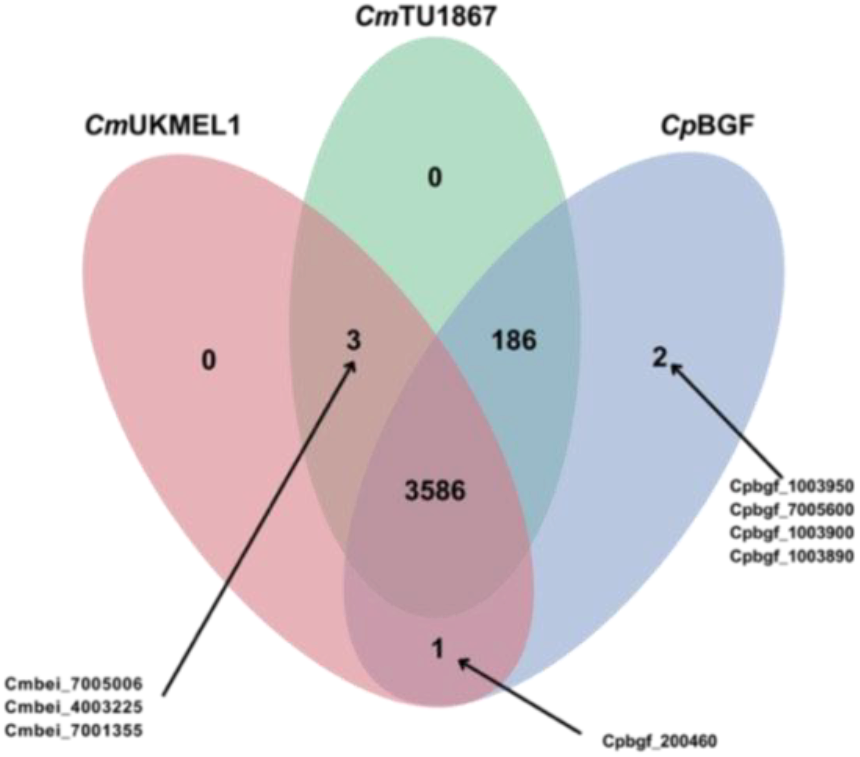
Venn diagram of ortholog search results following manual validation. Orthogroup comparison among the new *Cm*TU1867, the previous *Cm*UKMEL1, and the newly released reference genome, *Cp*BGF. See Figure 5 for the pre-validation results. Arrows link gene IDs to their orthogroups.

While annotating the genome, we noticed several genes that were annotated as a single long transcript in *Cm*UKMEL1 but as two distinct genes in *Cp*BGF. Upon investigation, we discovered that these gene annotations vary in size in several *Cryptosporidium* species. In *Cm*TU1867, the protein is annotated as one single long transcript for the 22 cases described in Table 2 since it is unlikely that the whole ORF could be annotated with no errors in the annotation software and since it is annotated as a single gene in *Cm*UKMEL1 and other *Cryptosporidium* spp. A lack of RNAseq evidence for *C. meleagridis* made it challenging to validate whether these genes exist as a single long gene in nature. We made a note that the gene is annotated as two or three distinct genes in other species in the annotation file for *Cm*TU1867 (two of the 20 proteins are annotated as 3 proteins in *Cp*BGF). Five *C. meleagridis* orthologs of *C. parvum* sub-telomeric genes could not be found in the current assembly including: cpbgf_1003890, cpbgf_1003900, cpbgf_1003950, and cpbgf_7005600 as well as cpbf_200460 which was observed in CmUKMEL but not CmTU1867 (Figure 3). These gene differences probably result from the incomplete *C. meleagridis* TU1867 sub-telomeric regions as we did not assemble many telomeres. However, it is also possible that those genes do not exist in *C. meleagridis*. This determination will require a *C. meleagridis* T2T assembly.

## Methods

### Whole Genome Sequencing and Assembly

*C. meleagridis* isolate TU1867 genomic DNA was obtained from BEI Resources (cat. number NR-2521 ATCC, Manassas, VA). A total of 10 ng of *C. meleagridis* DNA was amplified through whole genome amplification using multiple displacement amplification (MDA), followed by T7 endonuclease debranching yielding 400 ng debranched DNA following^13^ (Figure 4). ONT library preparation was performed using the SQK-RBK004 Rapid Barcoding Sequencing Kit (Oxford Nanopore Technologies, Oxford, UK) as per the manufacturer’s instructions. Sequencing was performed on an ONT MinION device with R9.4.1 flow cells and base called by guppy v.6.4.2 using the high-accuracy base call model.. The long-read fastq reads were assembled using Flye v.2.8.2^15^ with the --nano-raw option and -g 9m. The draft long read based genome was polished with PolyPolish v.0.5.0^16^ using default parameters to increase the accuracy of the base calls by using *C. meleagridis* strain TU1867 Illumina sequences (SRX253214) generated by others. Intermediate files needed for PolyPolish were generated using BWA v.0.7.17^17^. The resulting contigs were ordered and oriented to match the reference *Cp*BGF genome assembly using AGAT^18^ v. 1.1.0 PERL script agat_sq_reverse_complement.pl and GenomeTools^19^. Contig orientation was using the progressive Mauve alignment v 1.1.3^20^ in Geneious Prime v 2023.2.1^21^. Contamination was detected by searching the NCBI nr database using BLAST^22^ (blastx default parameters) and FCS-GX^23^. Contaminant contigs were removed from further analysis. We manually searched the contigs for telomeres. Telomeres were also identified, as in *Cp*BGF^24^ using the telomere-locating python script FindTelomeres to find the *Cryptosporidium* telomere repeat 5’-CCTAAA-3’ and its complement at the ends of assembled contigs (https://github.com/JanaSperschneider/FindTelomeres). The unassembled ONT long-reads were also searched for this telomere repeat with FindTelomeres and reads with telomeres were mapped back to the genome assembly using minimap2 (default parameters). Read-mapping to the whole genome was done using minimap2 v.2.26 with the option – secondary=no to prevent multi-mapping. Genome statistics were generated using GenomeTools v.1.6.2^19^ programs gt stat and gt seqstat. AGAT v.1.1.0^18^ PERL scripts agat_sq_stat_basic.pl and agat_sp_statistics.pl were used to generate statistical information with default parameters.

**Figure 4.**
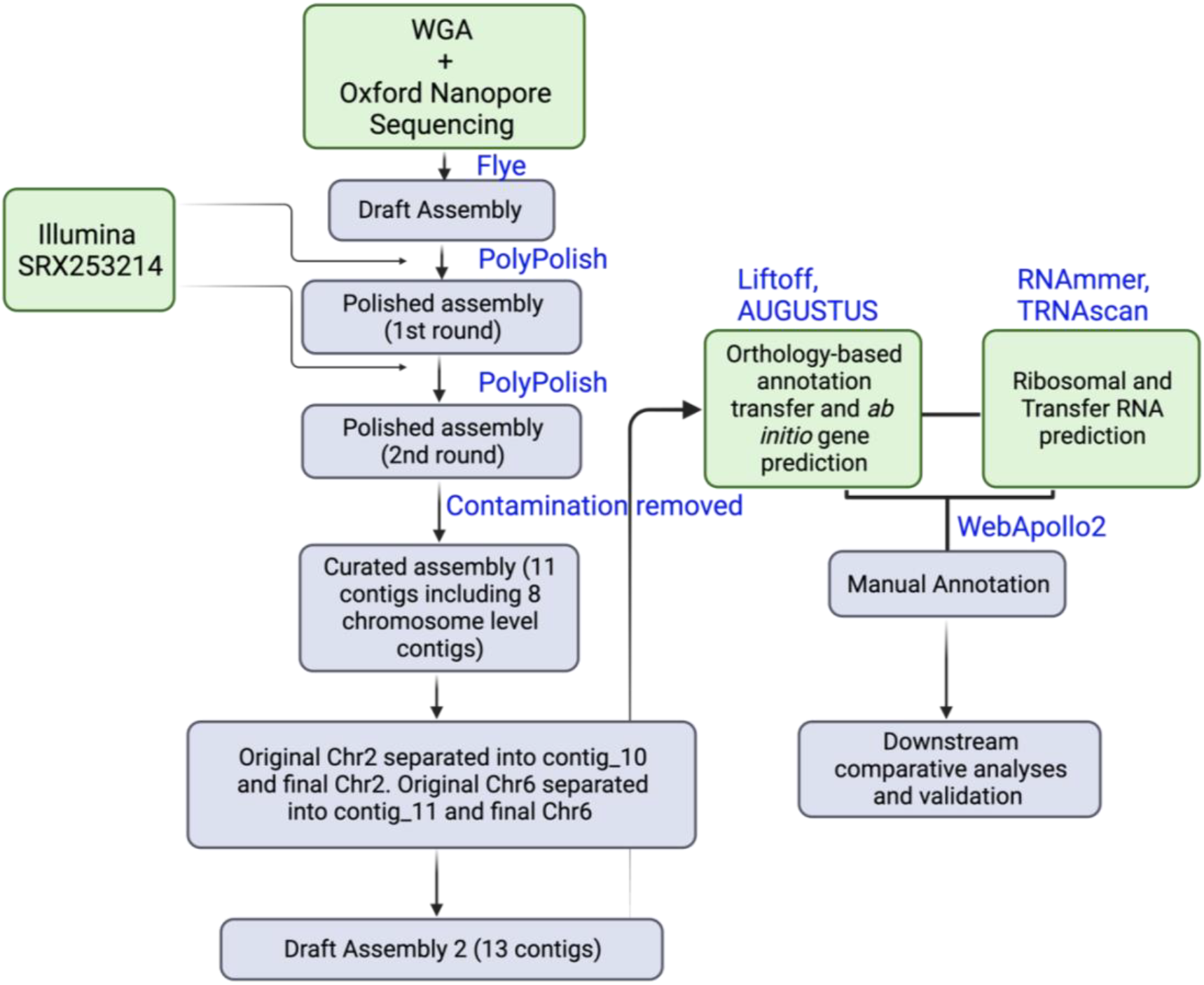
Experimental workflow for genome sequencing, assembly, annotation, and analysis. Bioinformatics workflow for assembly and annotation of the DNA derived from *Cm*TU1867 WGA. Green boxes represent main initial steps as well as new data used for parts of the pipeline and blue boxes represent subsequent downstream analyses of the data generated. Please refer to methods for additional details.

### Genome Annotation

Tracks for manual annotation were generated using a local Apollo2 server^25^ using two approaches: (1) an orthology based annotation transfer using the tool Liftoff^26^ and (2) an *ab initio* gene prediction using Augustus^27^ trained with *C. parvum* IOWA-ATCC and *Cm*UKMEL1 protein sequences from CryptoDBv.50 with the -copies flag to look for extra gene copies and otherwise default parameters. Annotation Liftoff tracks were created from the current *Cm*UKMEL1, *Cp*BGF, and *Cp*IOWA-ATCC annotated genes with the -copies flag to look for extra gene copies. In situations where AUGUSTUS and Liftoff gene structures disagreed, the conflicting gene models were searched in BLASTp in CryptoDB to check for the gene structure that was most abundant in existing annotations. As there is no available RNA-seq data for *C. meleagridis* there is no way to confirm gene predictions and UTRs are not annotated. Tracks for prediction and manual annotation of rRNAs were created using RNAmmer 1.2^14^ (S -euk, - m lsu, ssu). TRNAscan 2.0^28^ was used to predict tRNAs using default parameters. Functional annotation was generated with Blast2GO^29^ (using blastp, the nr database nr, word size 5, and e-value 1e-5) and compared with results from the reference telomere-to-telomere *Cp*BGF genome functional annotation. Edits to the *Cm*TU1867 gff file gene names were performed with basic bash and awk commands.

### Comparative Genomics

A comparison of orthologous genes between the new *C. meleagridis* assembly and the previous *C. meleagridis* assembly^30^ was completed using OrthoFinder v2.5.4^31^ with default parameters and visualized using OrthoVenn3^32^. Figure 3 represents the orthology results following extensive manual validation (Figure 5 and Table 3) of each orthogroup difference. Manual analyses utilized both NCBI BLASTp and CryptoDB^33^ BLASTp. Orthology, genome, and rRNA comparisons were created using Circos^34^, TBTools^35^, and JupiterPlot^36^. Comparisons of rRNA clusters in *Cp*BGF, *Cp*IOWA-ATCC, and *Cm*TU1867 were performed using RNAmmer^14^ with default parameters. *Cm*TU1867 long reads were mapped back to contig regions containing 5S rRNA clusters using minimap2 with --secondary=no to account for multi-mapping.

**Figure 5.**
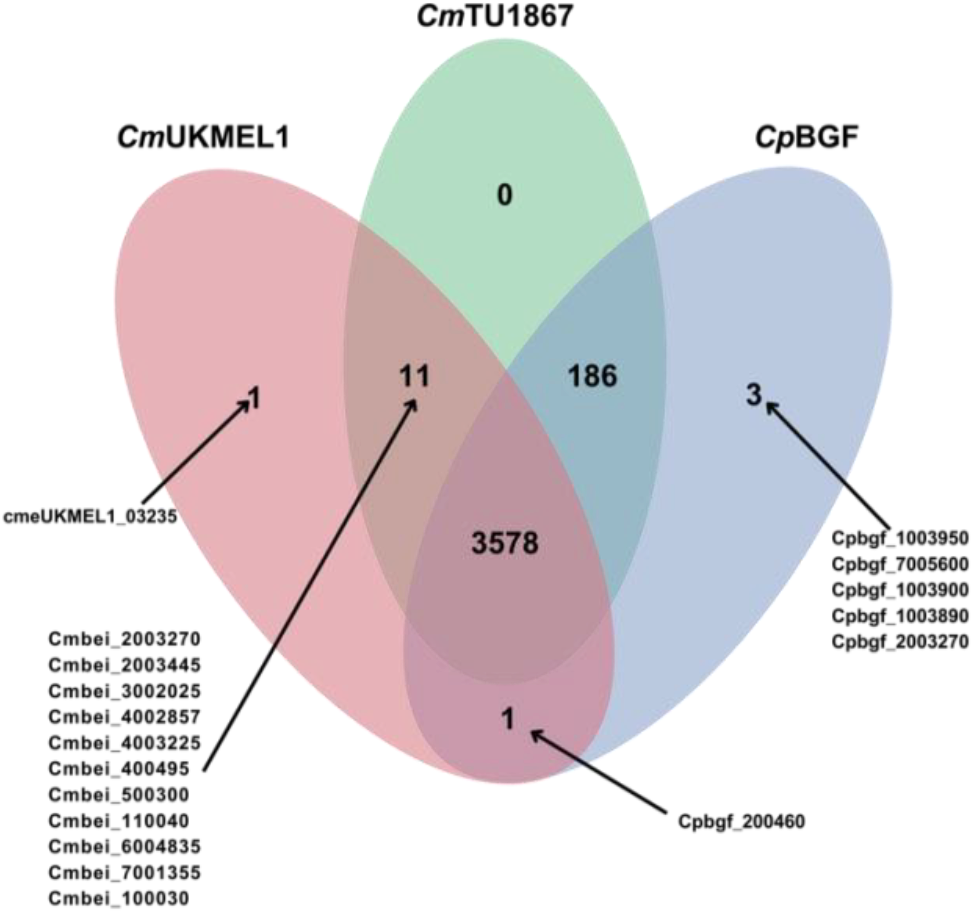
Ortholog search results shown in a Venn diagram. Orthogroup comparison among the new *Cm*TU1867, the previous *Cm*UKMEL1, and the newly released reference genome, *Cp*BGF prior to validation and correction. Arrows point to orthogroups containing the indicated gene IDs.

## Data Records

The genomic sequences, reads, and metadata for the *Cryptosporidium meleagridis* TU1867 strain have been deposited in the NCBI GenBank under BioProject PRJNA1022047. The sequence was polished with reads from NCBI GenBank SRA SRR793561. The fully assembled and annotated sequence was submitted to the NCBI GenBank under submission ID SUB14212942 and will be released as soon as it completes processing.

## Technical Validation

*Cm*TU1867 assembly completeness was evaluated using the Benchmarking Universal Single-Copy Orthologs (BUSCO) software v.5.5.0^37^ to search against apicomplexan databases (apicomplexa_odb10) which contain 446 orthologous single-copy genes in total. The results showed an overall completeness score of 96.6% (n=446). Of these, 430 (96.4%) single-copy genes were retrieved of which 1 (0.224%) was duplicated. These results indicate high completeness of the genome assembly.

Further analysis of the assembly and annotated protein encoding regions utilized an orthology comparison of *Cm*TU1867, *Cm*UKMEL1, and *Cp*BGF with the OrthoFinder algorithm in OrthoVenn3 (Figure 5). Orthogroups belonging to *Cm*UKMEL1 only, *Cp*BGF only, *Cm*UKMEL and *Cp*BGF only, and *Cm*UKMEL1 and *Cm*TU1867 only were extensively analyzed (Table 3). Several genes found only in *Cp*BGF were shown to be subtelomeric in both *Cm*UKMEL1 and *Cm*BEI and thus likely missing from the incomplete chromosome ends of CmTU1867. Several genes encoding short < 100 amino acid proteins found in both *Cm*BEI and *Cm*UKMEL1 exist in *Cp*BGF but are unannotated. Following these analyses, a new Venn diagram (Figure 3) was created that represents the revised, validated findings.

## Code Availability

Pipelines and code involved in processing the data were executed by following the respective manuals of the bioinformatics software programs used. No custom scripts were generated in this study.

## Acknowledgements

This work was funded by NIH R01AI14866 to JCK and TCG.

## Author contributions

JCK, RPB and TCG conceived the study; RPB, MSB, and LRP generated the genome assembly and LRP and RPB performed annotation; LRP performed analyses; LRP, RPB and JCK wrote the manuscript; TCG and JCK provided oversight and funding; All authors edited the manuscript.

